# Heightened NLRP3 inflammasome activation is associated with aging and CMML diseases severity

**DOI:** 10.1101/2022.01.27.477992

**Authors:** Nicola Andina, Louise de Meuron, Annatina Sarah Schnegg-Kaufmann, Camille Ansermet, Giuseppe Bomaci, Nino Keller, Naomi Porret, Anne Angelillo-Scherrer, Nicolas Bonadies, Ramanjaneyulu Allam

## Abstract

Aging causes chronic low-grade inflammation known as inflamm-aging. It is a risk factor for chronic myelomonocytic leukemia (CMML), a hematological malignancy that is most prevalent in older people. Recent studies suggest a critical role for the NLRP3 inflammasome in inflamm-aging. However, the mechanisms involved in NLRP3 activation in aging and its involvement in CMML progression are not fully understood. Here, we report that aging increases interleukin-1β production upon NLRP3 activation in human CD14^+^ monocytes. Interestingly, we found that Toll-like receptor (TLR) 1/2 agonist Pam3Cysk4 directly activates NLRP3 inflammasome without the requirement of second activation signal in monocytes from older but not from younger healthy donors. Further, we observed a dichotomous response to NLRP3 inflammasome activation in monocytes from CMML patients. Intriguingly, CMML patients with heightened NLRP3 activation showed increased treatment dependency and disease severity. Collectively, our results suggest that aging causes increased sensitivity to NLRP3 inflammasome activation at cellular level, which may explain increased inflammation and immune dysregulation in older individuals. Furthermore, NLRP3 inflammasome activation was dysregulated in CMML and positively correlated with disease severity.

## Introduction

As life expectancy of humans increases, chronic, as well as age related health problems have become a major issue for the healthcare system (Brown, 2015). One of the hallmarks of aging is the decline of the immune responses particularly adaptive immune system known as immunosenescence (Shaw et al., 2010; López-Otín et al., 2013), which explains the reduced response to a variety of immunogenic stressors such as infections, vaccines, and malignant cells in elderly people. Conversely, age-related changes in elderly people also lead to chronic low-grade inflammation with increase in pro-inflammatory cytokines such as interleukin-6 (IL-6) and interleukin-1β (IL-1β) and tumor necrosis factor (TNF) (Franceschi et al., 2000; Ferrucci and Fabbri, 2018). This chronic inflammatory state, collectively referred to as ‘inflamm-aging’, is a known risk factor for several non-communicable chronic disorders such as arthritis, atherosclerosis, diabetes, and cancer, including myeloid malignancies (Franceschi et al., 2018; Ferrucci and Fabbri, 2018). Several mechanisms can contribute to inflamm-aging, ranging from mitochondrial dysfunction, dysregulation in the ubiquitin-proteasome system, epigenetic alteration, increased activation of DNA damage response, defect in autophagy and changes in gut microbiome (Fülöp et al., 2019; Fulop et al., 2018). In addition, recent studies suggest a role for NLRP3 (NOD-, LRR- and pyrin domain-containing protein 3) inflammasome activation in inflamm-aging (Youm et al., 2013; Latz and Duewell, 2018; Sebastian-Valverde and Pasinetti, 2020).

NLRP3 inflammasome is a multiprotein signaling complex and its activation leads to caspase-1 mediated release of the pro-inflammatory cytokines IL-1β and IL-18, and initiates Gasdermin-D mediated cell death known as pyroptosis (Broz and Dixit, 2016). The classical NLRP3 inflammasome signaling is a two step-process that requires a priming (signal-1) and an activation signal (signal-2) (Swanson et al., 2019). During the initial priming step, stimulation of pattern recognition receptors (PRRs), such as Toll-like receptors (TLRs), lead to nuclear factor-κB (NF-κB) activation, which mediates transcriptional upregulation of the inflammasome components NLRP3 and pro-IL-1β. During the activation step, a signal induced by several pathogen-associated molecular patterns (PAMPs) or danger-associated molecular patterns (DAMPs) including bacterial toxins, RNA viruses, pathogenic crystals and altered cellular components, leads to NLRP3 activation and formation of an inflammasome complex (Gong et al., 2020; Henao-Mejia et al., 2014). In addition, an alternative NLRP3 inflammasome activation pathway has been recently described in human monocytes, wherein TLR4 stimulation by LPS leads to direct NLRP3 inflammasome activation independent of a second signal (Gaidt et al., 2016). This pathway is dependent on caspase-8 downstream of TLR4 and does not activate pyroptosis (Gaidt et al., 2016; Gaidt and Hornung, 2017). NLRP3 activation promotes chronic low-grade inflammation and is responsible for functional decline in aging (Latz and Duewell, 2018; Youm et al., 2013). However, mechanisms that contribute to NLRP3 inflammasome activation in aging are not completely understood. Though few studies showed decreased or unaltered NLRP3 activation in aged murine monocytes/macrophages (Bauernfeind et al., 2016; Ramirez et al., 2012; Stout-Delgado et al., 2012), whereas systematic and functional studies addressing NLRP3 inflammasome activation in healthy adult and aged human monocytes/macrophages are lacking.

Corroborating NLRP3 role in inflamm-aging, it is involved in several age-related disorders including chronic myelomonocytic leukemia (CMML) (Sebastian-Valverde and Pasinetti, 2020; Ratajczak et al., 2020; Duewell et al., 2010; Vandanmagsar et al., 2011; Henao-Mejia et al., 2014; Hamarsheh et al., 2020). Supporting this, chronic inflammatory diseases are frequently observed in patients with CMML (Solary and Itzykson, 2017; Kipfer et al., 2018).

CMML is a clonal myeloid neoplasm with risk of transforming into acute myeloid leukemia (AML) and eventually poor survival (Solary and Itzykson, 2017). CMML is nearly confined to elderly, the average patient diagnosed with CMML is 72 years old and the median survival ranges from less than 1 year to almost 3 years (Benzarti et al., 2019). According to the definition of the World Health Organization, diagnosis of CMML requires persistently elevated peripheral blood monocytes (≥1*10^9^/L), with monocytes accounting for at least 10% of the white blood cell count (WBC) (Arber et al., 2016). Recently, it has been shown that CMML patients with a *Kras^G12D^* mutation have increased NLRP3 inflammasome activation in peripheral blood mononuclear cells (PBMCs) compared to other CMML patients (Hamarsheh et al., 2020). However, it remains unknown if inflammasome mediated chronic inflammation is responsible for CMML disease progression.

In this study, we investigated the impact of aging on NLRP3 inflammasome activation using human CD14+ monocytes from young and older healthy donors. We found that aging increases NLRP3 inflammasome activation. Furthermore, we found a positive association for increased NLRP3 activation in monocytes and disease severity in CMML patients.

## Results

### Aging increases NLRP3 inflammasome activation in human monocytes

To understand whether aging contributes to dysregulation of NLRP3 inflammasome activation in humans, we studied inflammasome activation using monocytes from healthy donors (HD) of two different age groups: younger HD (30-35 years) and older HD (60-65 years). Monocytes are important innate immune cells, that produce inflammatory cytokines and are believed to contribute to inflamm-aging (Maeyer and Chambers, 2020). Monocytes represent approximately 10% of total circulating blood leukocytes in humans (Maeyer and Chambers, 2020). Immunophenotypically, these cells consist of classical (CD14^+^CD16^-^), non-classical (CD14^dim^CD16^+^) and intermediate (CD14^+^CD16^+^) populations (Ziegler-Heitbrock et al., 2010). Classical monocytes are most abundant, accounting for 90-95% of all peripheral blood monocytes (Mildner et al., 2013). We isolated CD14^+^ monocytes from peripheral blood, which covers more than 95% of total monocytes. We checked their purity and morphology by FACS and cytospin (**Fig.1 A and B**). Next, we primed these monocytes with TLR2 ligand Pam3CSK4 and stimulated with nigericin to induce canonical NLRP3 inflammasome activation. We also stimulated cells with LPS alone to induce alternative NLRP3 inflammasome activation (Gaidt et al., 2016). Interestingly, in contrast to immunosenescence phenotype, monocytes from older HD showed increased mature IL-1β production for the canonical and alternative NLRP3 agonists (**Fig. 1C**). Confirming above results, monocytes from older HD showed increased IL-1β and caspase-1 cleavage compared to younger HD on western blot (**Fig. D**). These results suggest that aging increases sensitivity for NLRP3 inflammasome activation.

**Figure 1.**
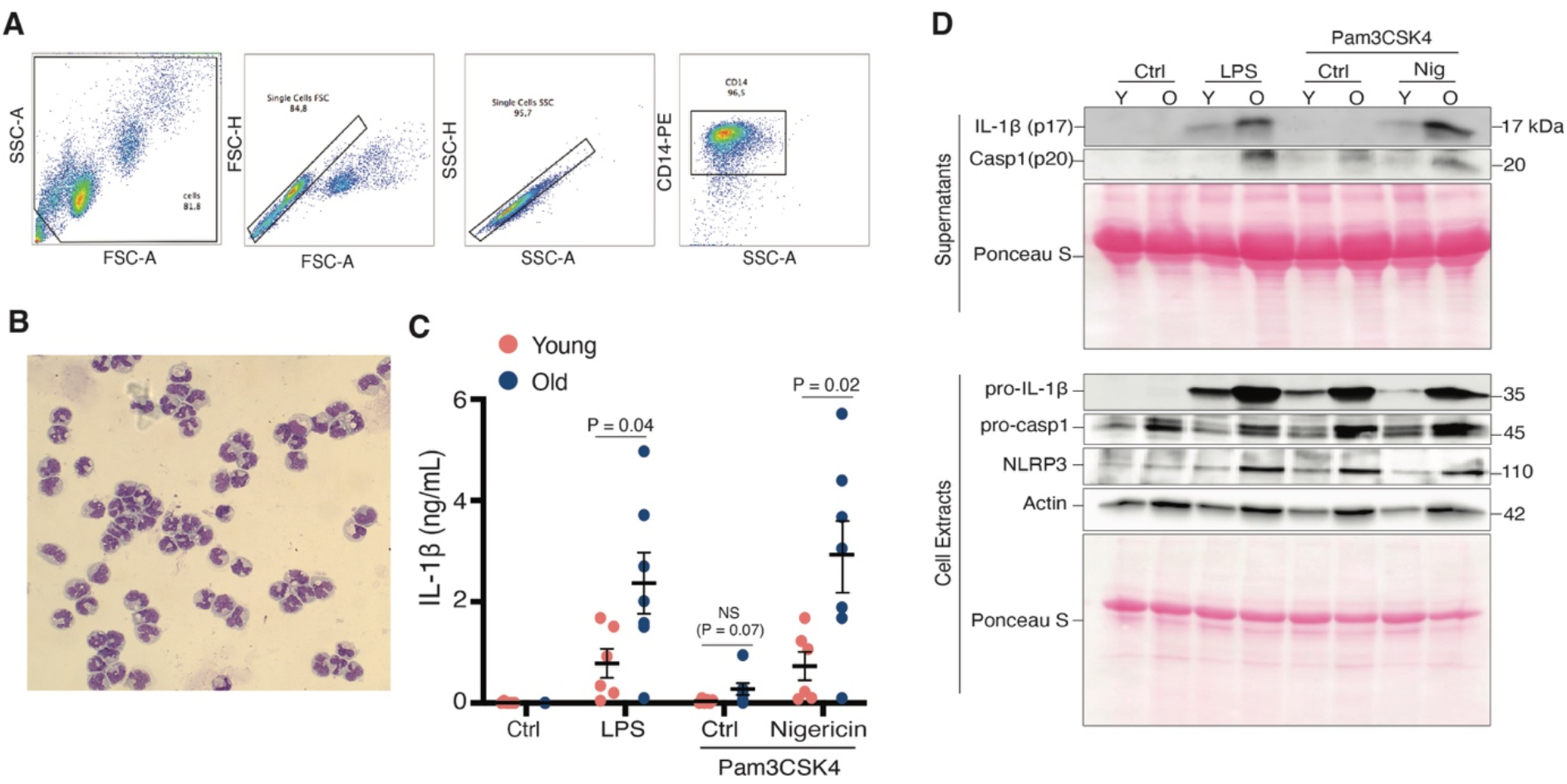
Aging increases NLRP3 inflammasome activation. CD14^+^ monocytes were isolated from healthy donors of two different age groups: younger healthy donors (30 - 35 years of age; Y, young) and older healthy donors (60 - 65 years of age; O, old). (**A and B**) FACS analysis (A) and cytospin (B) was performed to check monocyte purity and morphology. (**C and D**) Monocytes were stimulated with LPS (2 μg/mL) or Pam3CSK4 (2 μg/mL) for 16 hours and stimulated for 2 hours with or without 5 μM of nigericin (Nig). Supernatants were collected and IL-1β ELISA was performed (C). Data shown as mean ± SEM. Statistical analyses were performed using a two-tailed *t*-test. Cell lysates and supernatants were analyzed for pro- and cleaved-forms of caspase-1 and IL-1β by western blot (D). Blots are representative of three independent experiments.

### Aging amplifies inflammasome priming via increased NF-κB activation

As aforementioned, activation of canonical NLRP3 inflammasome requires a priming and an activation signal (Swanson et al., 2019). Upon priming, we observed increased pro-IL-1β levels in older HD, suggesting that with age inflammasome priming is amplified (**Fig. 1D**). To understand the involved mechanism, monocytes from younger and older HD were stimulated with the Pam3CSK4 and analyzed for canonical and non-canonical NF-κB signaling. Monocytes from older HD displayed decreased levels of the IκBα and increased levels of NF-κB2 (p52) (**Fig. 2A**), suggesting that aging can increase both canonical and non-canonical NF-κB signaling, respectively. To confirm NF-κB activation, we monitored the production of pro-inflammatory cytokine TNF after LPS and Pam3CSK4 stimulation and found that monocytes from older HD exhibited a trend of enhanced secretion of TNF compared to younger HD (**Fig. 2B**). To further ascertain this, we inhibited NF-κB signaling and NLRP3 activation by dexamethasone and MCC950, respectively (Auphan et al., 1995; Coll et al., 2015). Dexamethasone treatment significantly reduced IL-1β and TNF levels but MCC950 only inhibited IL-1β in monocytes from younger and older HD (**Fig. 2C and D**). Collectively, these results suggest that aging amplifies inflammasome priming potentially via increased NF-κB activation.

**Figure 2.**
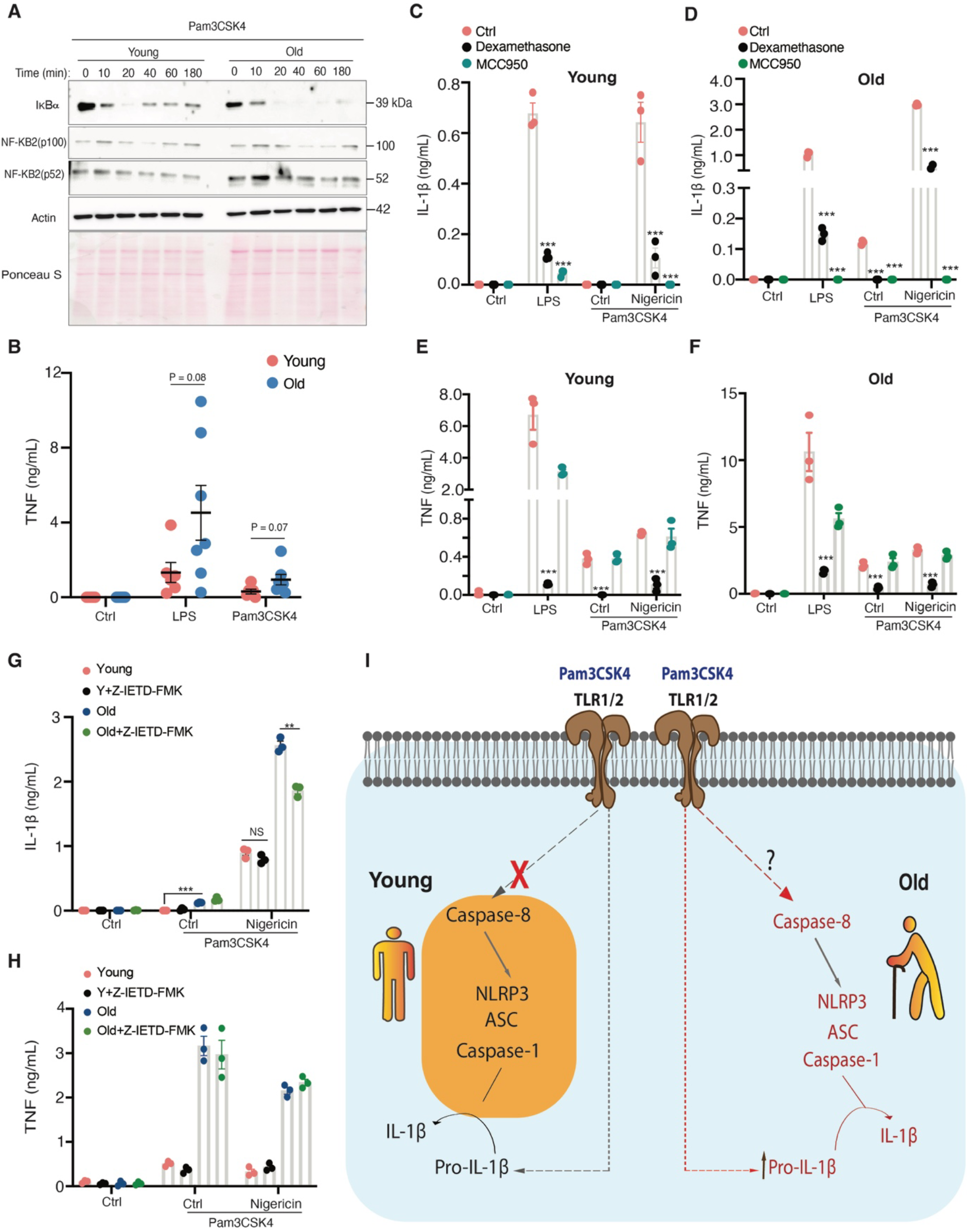
Aging increases NF-κB activation and sensitivity for TLR1/2 ligand. **(A)** Monocytes from younger and older healthy donors were stimulated with Pam3CSK4 (1 μg/mL) for different time points as indicated. Total cell lysates were analyzed for NF-κB activation by western blot with indicated antibodies. Blots are representative of two independent experiments. (**B)** Monocytes from younger and older healthy donors were stimulated with Pam3CSK4 (2 μg/mL) or LPS (2 μg/mL) for 18 hours. Supernatants were analyzed for TNF by ELISA. Data is shown as mean ± SEM. Statistical analyses were performed using a two-tailed *t*-test. (**C-F**) Monocytes from younger and older healthy donors were pre-treated with or without dexamethasone (100 nM; NF-κB inhibitor) or MCC950 (2 μM; NLRP3 inhibitor) for 1 hour. After, cells were stimulated with LPS (2 μg/mL) or Pam3CSK4 (2 μg/mL) for 16 hours and with or without 5 μM of nigericin for 2 hours. Supernatants were analyzed for IL-1β (C, D) and TNF (E, F) by ELISA. Data is shown as mean ± SEM. Statistical analyses were performed using a two-tailed *t*-test. (**G and H**) Monocytes from younger and older healthy donors were pre-treated with or without Z-IETD FMK (4 μM; caspase-8 inhibitor) for 1 hour. Later cells were stimulated with LPS (2 μg/mL) or Pam3CSK4 (2 μg/mL) for 16 hours and stimulated for 2 hours with or without 5 μM of nigericin. Supernatants were analyzed for IL-1β (G) and TNF (H) by ELISA. Data is shown as mean ± SEM. Statistical analyses were performed using a two-tailed *t*-test. (**I**) Schematic figure showing that Pam3CSK4 directly activates NLRP3 inflammasome in monocytes from older but not from younger healthy donors. This activation is akin to LPS inducing the alternative NLRP3 inflammasome might depend on caspase 8.

### Aging licenses TLR1/2 signaling to activate alternative NLRP3 inflammasome activation

In human myocytes LPS directly activates NLRP3 inflammasome and produce IL-1 β in the absence of a secondary activating signaling known as alternative NLRP3 inflammasome activation (Gaidt and Hornung, 2017). As expected, LPS directly activated IL-1 β production in the absence of a second signal in monocytes from younger and older HD (**Fig 1C, Fig 2C and D**). Interestingly, akin to LPS, TLR1/2 ligand Pam3CSK4 directly induced IL-1β production in monocytes from older but not from younger HD (**Fig 1C**). Although the increase of IL-1 β production in monocytes from older HD is not statistically significant, there is an increased trend. Further, in monocytes from older HD, NLRP3 inflammasome inhibitor MCC950 completely abolished Pam3CSK4 induced IL-1β production, suggesting this activation is dependent on NLRP3 (**Fig 2D**). To check whether caspase-8 is involved in Pam3CSK4 induced IL-1β production akin to LPS, we pre-treated the cells with caspase-8 inhibitor Z-IETD FMK and stimulated with Pam3CSK4. Although, caspase-8 inhibitor Z-IETD FMK did not show inhibitory effect on Pam3CSK4 induced IL-1β production, it inhibited Pam3CSK4 and nigericin induced IL-1β production in monocytes from older HD but not in monocytes from younger HD (**Fig 2G and H**). This indicates that Pam3CSK4 can potently activate NLRP3 activation through caspase-8 in monocytes from older HD. Inhibiting caspae-8 alone can lead to receptor-interacting kinase 3 (RIP3) dependent necroptosis (Kaiser et al., 2011) and might interfere with inhibitory effector of Z-IETD FMK. So, further studies are needed to rule out caspase-8 in Pam3CSK4 induced IL-1β production. Altogether, these results suggest that aging increases sensitivity to Pam3CSK4 via the alternative NLRP3 inflammasome activation pathway in monocytes (**Fig 2I**).

### Dichotomous NLRP3 inflammasome response in monocytes from CMML patients

CMML disease is generally affects elderly people and is characterized by elevated monocyte counts (monocytosis) (Solary and Itzykson, 2017; Benzarti et al., 2019). CMML shows frequent association (in 10 – 30% of cases) with systemic autoimmune and inflammatory diseases (SAIDs) (Grignano et al., 2016; Solary and Itzykson, 2017; Kipfer et al., 2018). SAIDs can either precede or follow the diagnosis of CMML. It is however unknown if SAIDs impact CMML prognosis. Since our results suggest that aging increases NLRP3 inflammasome activation, we investigated whether monocytes from CMML patients have increased NLRP3 inflammasome activation. Median age of CMML patients was 70.2 years (range 57.6 - 80.4 years) at the time of sample collection. Additional clinical characteristics are summarized in **Table 1**. Strikingly, we identified dichotomy in IL-1β production in response to Pam3CSK4/nigericin and LPS induced canonical and alternative NLRP3 inflammasome activation, respectively (**Fig. 3A**). In comparison to monocytes from older HD (median age 62.5 years), some CMML patients monocytes produced high levels of IL-1β (high responders, CMML-HR) and some produced low levels of IL-1β (low responders, CMML-LR) (**Fig. 3A**). Interestingly, this dichotomy was not observed for TNF production, where cytokine levels were comparable between monocytes from CMML patients and monocytes from older HD (**Fig. 3B**). Confirming above results, monocytes from CMML-HR or CMML-LR showed increased or decreased IL-1β and caspase-1 cleavage, respectively, compared to monocytes from older HD (**Fig. 3C**). To better understand the increased NLRP3 inflammasome activation in CMML-HR, we performed NLRP3 inflammasome activation in the presence of pathway inhibitors. As expected, dexamethasone pre-treatment significantly reduced IL-1 β and TNF production but MCC950 only inhibited IL-1β production (**Fig. 3D and E**). Further, akin to monocytes from older HD, Pam3CSK4 directly induced IL-1β production in CMML-HR responders and this response was inhibited by all three inhibitors (**Fig. 3D**). Interestingly, in these experiments caspase-8 inhibitor Z-IETD FMK significantly inhibited Pam3CSK4 induced IL-1β production in monocytes from CMML-HR (**Fig. 3D**). These results substantiate the above conclusion that caspase-8 is involved in Pam3CSK4 induced alternative NLRP3 inflammasome activation. Collectively, these results suggest increased canonical and alternative NLRP3 inflammasome activation in CMML-HR patients.

**Figure 3.**
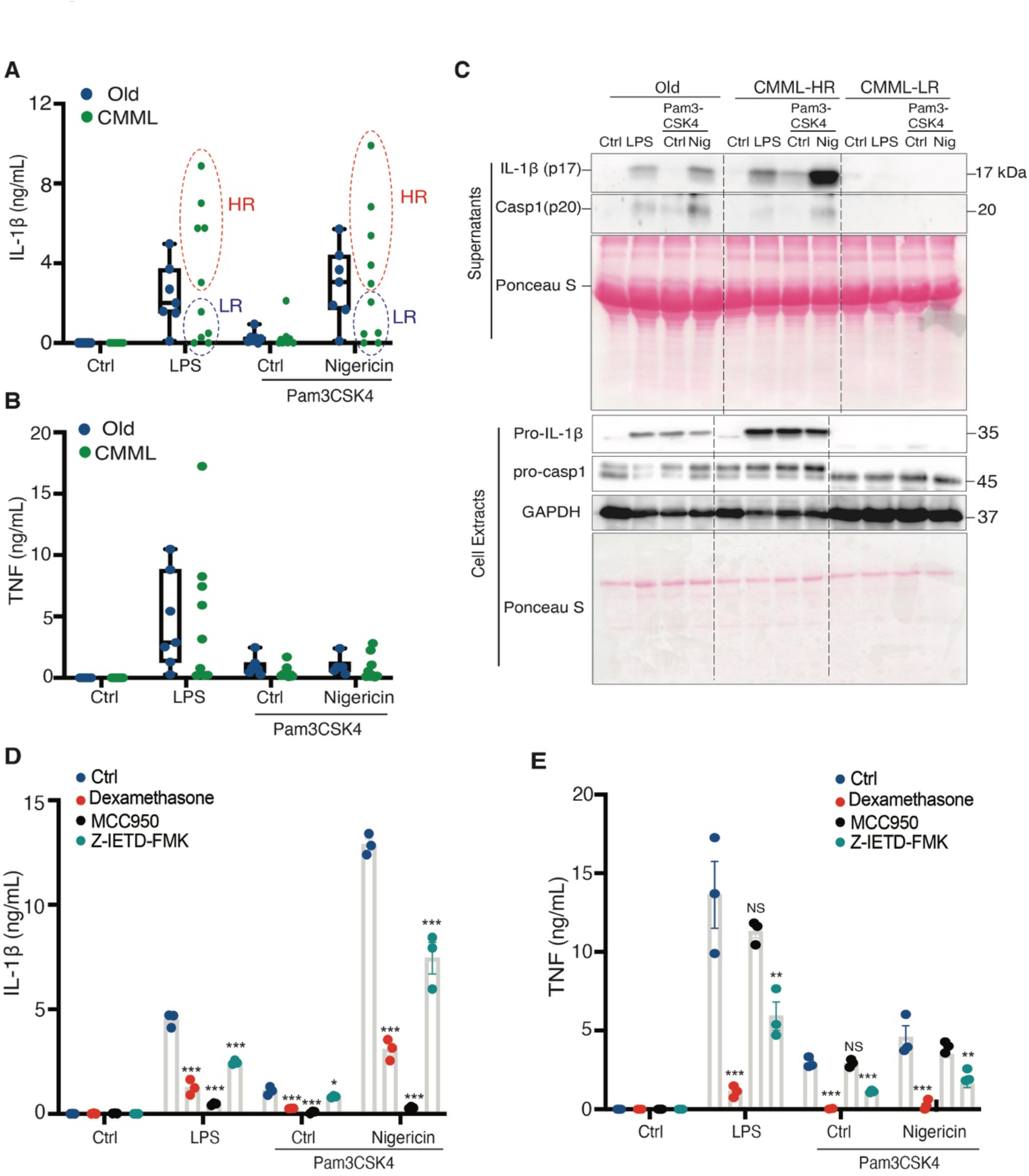
Dysregulated NLRP3 inflammasome activation in CMML monocytes. (**A-C**) Monocytes from older healthy donors (60 - 65 years of age, OHD) and CMML patients were stimulated with LPS (2 μg/mL) or Pam3CSK4 (2 μg/mL) for 16 hours and with or without 5 μM nigericin for 2 hours. Supernatants were analyzed for IL-1β (A) and TNF (B) by ELISA. Data is shown as mean ± SEM. Statistical analysis was performed using a two-tailed *t*-test. Cell lysates and supernatants were analyzed for pro- and cleaved-forms of caspase-1 and IL-1β by western blot (C). Blots are representative of three independent experiments. (**D and E**) Monocytes from older healthy donors and CMML patients were pre-treated with or without dexamethasone (100 nM; NF-κB inhibitor) or MCC950 (2 μM; NLRP3 inhibitor) or Z-IETD FMK (4 μM; caspase-8 inhibitor) for 1 hour. After, cells were stimulated with LPS (2 μg/mL) or Pam3CSK4 (2 μg/mL) for 16 hours and with or without 5 μM of nigericin for 2 hours. Supernatants were analyzed for IL-1β (D) and TNF (E) by ELISA. Data is shown as mean ± SEM. Statistical analysis was performed using a two-tailed *t*-test.

Differences in IL-1β production between CMML-LR and CMML-HR cannot be attributed to differences in age, as the median age at the time of monocyte isolation was similar (69.7 years for CMML-LR, 70.7 years for CMML-HR). Furthermore, age alone is unlikely to explain the difference in IL-1β production of monocytes from CMML patients compared to older HD: First, the difference in median age (7.7 years between older HD and CMML) at the time of monocyte isolation is relatively small, and CMML-LR patients were not consistently younger than older HD with comparatively higher monocytic IL-1β production. (**Fig. 3A**). Second, although there was a clear trend to higher TNF levels in response to LPS in monocytes from older compared to younger HD (**Fig. 3B**), no such trend was observed between monocytes from CMML patients compared to older HD (**Fig. 3B**), further underscoring that a separate, age-independent mechanism leads to increased monocytic NLRP3 inflammasome activation in a subset of CMML patients.

### NLRP3 inflammasome response associates with CMML disease severity

Next, we checked whether the observed difference in NLRP3 inflammasome activation was associated with CMML disease severity. Based on peripheral blood and bone marrow blast cell percentage CMML is classified into three different categories, with worsening prognosis: CMML-0, CMML-1, and CMML-2 (Onida et al., 2002; Schuler et al., 2014; Orazi and Germing, 2008; SH et al., 2017). Interestingly, CMML-LR entirely classified as CMML-0, whereas CMML-HR mainly classified as higher stage CMML (CMML-1, CMML-2) (**Table. 1**). This was also reflected by a trend to higher risk scores, namely the international prognostic scoring system (IPSS) (Greenberg et al., 1997) the revised IPSS (IPSS-R) (Greenberg et al., 2012) as well as the more CMML specific CMML-specific prognostic scoring system (CPSS) (Such et al., 2013), and the CPSS molecular score (Elena et al., 2016) (**Table. 1**). Accordingly, high risk mutations, such as *ASXL1*, *NRAS*, *RUNX1*, *SETBP1*, were more frequent in the CMML-HR group. The most frequently observed mutation in CMML are *TET2* mutations (around 45%) (Patnaik et al., 2016). *TET2* mutated cases were more likely to be in CMML-0 and are less likely to have low hemoglobin levels compared to *TET2* unmutated cases. Overall and in the absence of an additional *ASXL1* mutation, *TET2* mutations were associated with better outcome (Patnaik et al., 2016). Indeed, *TET2* mutations were present in 4 CMML-LR patients (NGS was performed for 4 out of 5 CMML-LR patients). On the other hand, only one patient had a *TET2* mutation in the CMML-HR group (NGS was performed for 3 out of 5 CMML-HR patients). In line with this, we observed shorter time to therapy (**Fig. 4A**) and decreased overall survival (**Fig. 4B**) in the CMML-HR group. Over the course of a follow-up of approximately two years all CMML-HR patients were eventually given CMML related therapy, which included the hypomethylating agent azacitidine, hydroxyurea, and erythrocyte transfusions, with or without application of erythropoietin. During a similar follow-up period, none of the CMML-LR patients received CMML related therapy, neither before monocytes were isolated, nor during follow-up time. Appropriately and in line with the large proportion of *TET2* mutations found in this group, hemoglobin levels were significantly higher in the CMML-LR patients (**Fig. 4C**) and negatively correlated to NLRP3 inflammasome response in monocytes (**Fig. S1**). There was however no significant difference in the white blood cell count (WBC) (**Fig. 4D**), the platelet count (PLT) (**Fig. 4E**), and the monocyte count (MONO) (**Fig. 4F**). We also did not observe any significant correlation between these cell counts and NLRP3 inflammasome response in monocytes (**Fig. S1**). Another clinical feature associated with inferior prognosis is the presence of palpable splenomegaly, which is seen in up to 30% of CMML patients (Hoversten et al., 2018). While splenomegaly was absent in 4 out of 5 CMML-LR patients, splenomegaly was present in all CMML-HR patients, except for one that was previously splenectomized because of an accident. Furthermore, in line with increased monocytic inflammasome response, we saw a trend to higher levels of c-reactive protein (CRP), and therefore more inflammation, in CMML-HR patients (**Fig. 4G**). However, and in contradiction to higher CRP levels, potentially CMML associated SAIDs were more abundantly reported in the CMML-LR group (**Table. 1**). Surprisingly, we also did not observe any significant correlation between CRP levels and NLRP3 inflammasome response in monocytes (**Fig. S1**). Taken together, in our small cohort, monocytic NLRP3 inflammasome response was associated with CMML disease severity.

**Figure 4.**
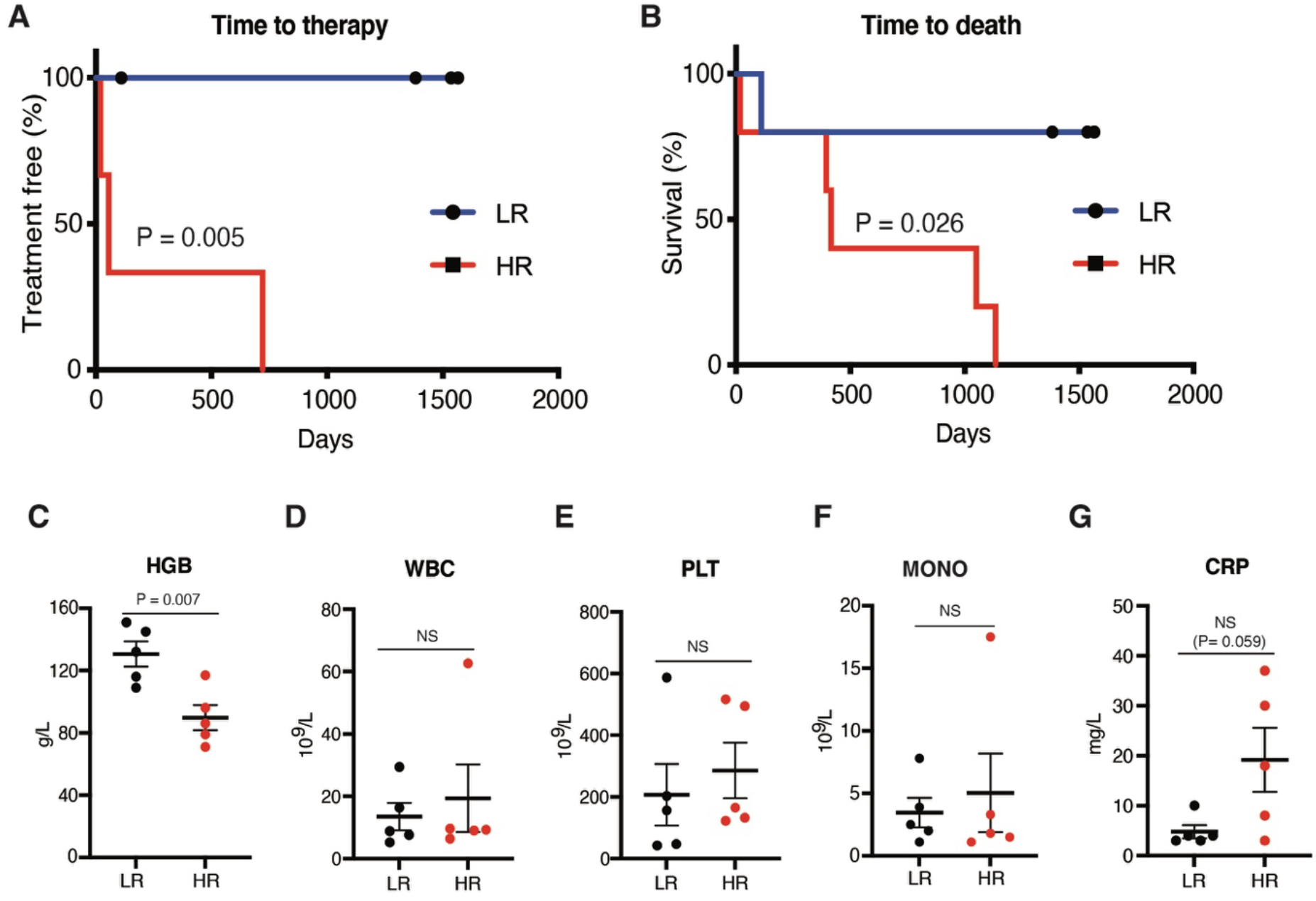
Association of NLRP3 inflammasome response with CMML clinical and laboratory parameters. Based on their NLRP3 inflammasome response, CMML patients were either grouped in the low responder group (LR) or in the high responder group (HR). (**A and B**) Kaplan-Meier curves for time to therapy (A), and time to death (B). Starting point for both was the time of monocyte isolation. This was done with a median of 405 days (0 – 3582 days) after the diagnosis of CMML was made. (**C-G**) Hemoglobin levels (HGB, C), white blood cell count (WBC, D), platelet count (PLT, E), monocyte count (MONO, F), and c-reactive protein levels (CRP, G) in CMML patients at the time of monocyte isolation. Data is shown as mean ± SEM. Statistical analysis was performed using a two-tailed *t*-test.

## Discussion

In this study, we demonstrated that aging causes dysregulation of NLRP3 inflammasome activation in human monocytes. Previous studies regarding the aging impact of TLR stimulation on monocyte activation have resulted in conflicting results (Shaw et al., 2010; Hearps et al., 2012; Shaw et al., 2013; Duin et al., 2007). Disparate outcomes could be dependent on several factors including use of total PBMCs versus pure monocyte population, various experimental protocols, and different enrolment requirements for individual subject selection. However, to our knowledge there are no studies that directly addressed aging related NLRP3 inflammasome activation in human monocytes. Our *ex vivo* experiments with primary human monocytes demonstrate increased NLRP3 inflammasome activation in monocytes from older healthy blood donors. These results are in line with growing evidence of NLRP3 inflammasome activation in age related diseases (Sebastian-Valverde and Pasinetti, 2020; Ratajczak et al., 2020; Duewell et al., 2010; Vandanmagsar et al., 2011; Henao-Mejia et al., 2014; Hamarsheh et al., 2020). How NLRP3 inflammasome is activated in older individuals is not completely understood. One possibility is accumulation of sterile activators with age such as DAMPs and ROS. Theses activators can trigger NLRP3 inflammasome activation in monocytes and contributes to inflamm-aging. In addition, our results also reveal that activation of the alternative NLRP3 inflammasome pathway by TLR1/2 ligand Pam3CSK4 takes place in monocytes from older but not from younger individuals. Our results suggest that aging has a cell-intrinsic effect on monocytes that renders them sensitive for Pam3CSK4 activation. This could explain why older individuals have increased inflammatory reactions during infection compared to younger individuals (Kale and Yende, 2011). How aging leads to these pro-inflammatory cell intrinsic changes is unknown. It is possible that aging mediated mitochondrial dysfunction, epigenetic alteration and defect in autophagy contributes to these changes. However, further studies are needed to understand the mechanism that contributes to cell intrinsic changes for inflammatory signals in monocytes from older individuals. Further, it will be interesting to study whether DAMPs that accumulate with age also activate monocytes from older individuals akin to Pam3CSK4.

CMML is a disorder of older individuals, and SAIDs are frequently observed in patients with CMML (Grignano et al., 2016; Solary and Itzykson, 2017; Kunnumpurath et al., 2020). However, it remains unknown if chronic inflammation is responsible for CMML or CMML promotes chronic inflammation. In this study, we have shown dysregulation of inflammasome activation in CMML patients. We found that monocytes from CMML patients had a dichotomous response for NLRP3 inflammasome activation, with and high and low responders. Furthermore, NLRP3 inflammasome response is positively associated with CMML disease severity according to IPSS, IPSS-R and CPSS. In line with this, a recent study shows that a *Kras^G12D^* mutation in mouse hematopoietic stem cells leads to a reminiscent of CMML phenotype, which is reversed by deleting the *Nlrp3* gene in this mouse (Hamarsheh et al., 2020). Additionally, they also found increased NLRP3 inflammasome activation in CMML patients with *KRAS^G12D^* mutation compared to patients with non-*KRAS^G12D^* mutations (Hamarsheh et al., 2020). In our small cohort, we did not find a *KRAS* mutation. This is not surprising as it is found only in low frequencies in CMML (3% in one report) (Grignano et al., 2016; Solary and Itzykson, 2017). We did however find that the presence of a *TET2* mutation was associated with lower NLRP3 inflammasome response and better outcome. Although, previous studies indicated that loss of *TET2* in mouse models increases inflammasome activation and contributes to atherosclerosis and insulin resistance (Fuster et al., 2017; Sano et al., 2018; Fuster et al., 2020), how *TET2* mutations alter inflammasome response in human monocytes needs to be examined. Further studies with a larger number of patient samples are required to clarify the correlation between inflammasome activation and disease severity in CMML patients and whether modulation of inflammasome activity may influence disease course.

Collectively, our exploratory study suggests that aging causes cell intrinsic changes in monocytes which contribute to increased NLRP3 inflammasome activation. These changes could be responsible for inflamm-aging and potentially involved in age-related diseases. Monocytic NLRP3 inflammasome activation is furthermore dysregulated in CMML, where it is positively associated with disease severity. Identifying cell-intrinsic changes and related molecular mechanisms could potentially lead to novel approaches to target age-related diseases, such as CMML.

## Acknowledgements

This work was supported by the Swiss National Science Foundation (PP00P3_183721 & PP00P3_190073), Novartis Foundation for Medical-Biological Research, and Olga Mayenfisch Stiftung to RA.

## Author contributions

NA, LM, ASK, CA, and GB performed experimental work. NB provided patients’ samples and data through the Swiss MDS Registry and Biobank platform. AAS helped with reagents and advice for experiments. RA and NA designed and supervised the research work. RA and NA wrote the manuscript with inputs from all other authors.

## MATERIALS AND METHODS

### Samples and ethics approval

Blood samples from healthy donors were obtained from the Interregional Blood Transfusion SRC, Bern, Switzerland. CD14^+^ monocytes were isolated from healthy donors of two different age groups: 30 - 35 years of age (referred to as “young healthy donors”) and 60 - 65 years of age (referred to as “old healthy donors”). Blood samples from CMML patients were obtained from the Department of Hematology, Inselspital, Bern and provided their informed consest to the Swiss MDS Registry/Biobank and further biomarker studies approved by the competent local ethics committee (2016-01917, 2017-02299, 2017-00699) The diagnosis was based on the criteria defined by the WHO in 2016/2017 (Orazi and Germing, 2008; SH et al., 2017; Choi and O’Malley, 2018).

### Isolation of monocytes from healthy and CMML patients

Monocyte isolation was carried out by using Anti-CD14^+^ Micro Beads (catalog 130-050-201, Miltenyi Biotec, Germany) from PBMCs separated from the blood samples. Isolated monocytes were analyzed by FACS using an anti CD14^+^ antibody (catalog 130-113-147, Miltenyi Biotec, Germany) to check their purity, and by cytospin to check their morphology and purity.

### Reagents

We purchased nigericin (N7143) and dexamethasone from Sigma-Aldrich, ultrapure LPS (LPS-EK), and Pam3Cysk4 (tlrl-pms) from Invivogen, Z-VAD-FMK (ALX-260-020) from Enzolifesciences, and MCC950 from Adipogen.

### Stimulation Experiment

Primary monocytes were cultured in RPMI 1640 GlutaMAX™-I medium (Invitrogen) supplemented with 2% (vol/vol) FBS (Amimed), 1% of penicillin and streptomycin (PAA Laboratories), at 37°C and 5% CO2. All cells were stimulated at a density of 1×10^6^ cells per mL. Monocytes were treated with LPS (2 μg/mL) or Pam3CSK4 (2 μg/mL) for 16 hours and stimulated for 2 hours with or without 5 μM of nigericin. Later, cell-free supernatants were analyzed by ELISA for cytokine secretion and cell pellets were lysed for western blot analysis.

### NF-κB activation

Monocytes were stimulated with TLR2 agonist Pam3CSK4 (1 μg/mL) at indicated time points. After stimulation cell lysates were isolated and western blot was performed.

### Western blot analysis

Precipitated media supernatants or cell extracts were analyzed by standard western blot technique. The following antibodies were used: Anti-human IL-1β antibody (12242), anti-NFκB2 (p100), anti-IκBa, anti-Phospho-IκBa, from Cell signaling. Anti-human caspase-1 (p20, Bally-1), and anti-NLRP3 (Cryo-2), from Adipogen. Anti-β-actin from Abcam.

### Cytokine measurement

Cell supernatants were analyzed for IL-1β and TNF cytokine secretion by ELISA according to the manufacturer’s instructions (eBioscience).

### Next generation sequencing (NGS)

NGS was performed on all CMML patient samples using the Illumina TruSight Oncology 500 assay. 523 tumor associated genes were analyzed, including known myeloid driver genes, such as: ABL1, ASXL1, BCOR, BRAF, CALR, CBL, CEBPA, CSF3R, DNMT3A, ETV6, EZH2, FLT3, GATA2, HRAS, IDH1, IDH2, IKZF1, JAK2, KIT, KRAS, MPL, MYD88, NF1, NPM1, NRAS, PHF6, PRPF8, PTPN11, RB1, RUNX1, SETBP1, SF3B1, SH2B3, SRSF2 (Exon 1), STAG2, TET2, TP53, U2AF1, WT1, *ZRSR2*.

### Statistical analysis

Data were expressed as mean ± SEM. Comparison between two groups was performed by two-tailed Etest. For Kaplan-Meier curves Log-rank (Mantel-Cox) test was used to test significant. For correlation analysis, we used Pearson’s correlation and significant was tested using twotailed *t*-test. A value of p<0.05 was considered statistically significant. All statistical analyses were calculated using Graph Pad Prism version 9.

