## Supplemental Figure 1 for "Heightened NLRP3 inflammasome activation is associated with aging and CMML diseases severity"

**Figure S1**

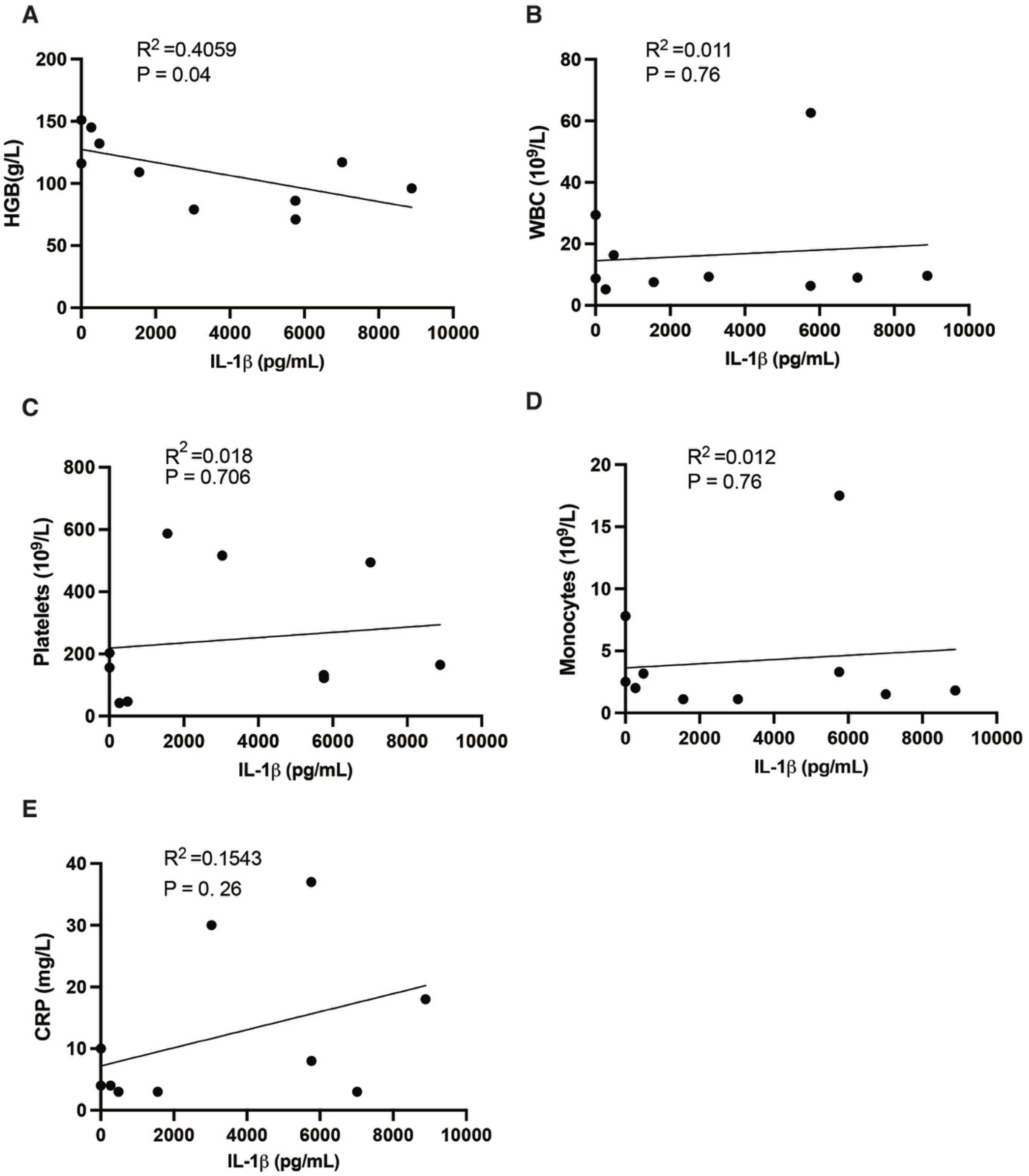


**Figure S1. Correlation of NLRP3 inflammasome response with CMML laboratory parameters**

(**A-E**) IL-1β levels from LPS induced CMML patient's monocytes were correlated with laboratory parameters. Hemoglobin (HGB, A), white blood cell count (WBC, B), platelet count (C), monocyte count (D), and c-reactive protein (CRP, E). For correlation analysis, we used

Pearson's correlation and significant was tested using two-tailed *t*-test. A value of $p<0.05$ was considered statistically significant.
